# Leveraging allelic heterogeneity to increase power of association testing

**DOI:** 10.1101/498360

**Authors:** Farhad Hormozdiari, Junghyun Jung, Eleazar Eskin, Jong Wha J. Joo

## Abstract

The standard genome-wide association studies (GWAS) detects an association between a single variant and a phenotype of interest. Recently, several studies reported that at many risk loci, there may exist multiple causal variants. For a locus with multiple causal variants with small effect sizes, the standard association test is underpowered to detect the associations. Alternatively, an approach considering effects of multiple variants simultaneously may increase statistical power by leveraging effects of multiple causal variants. In this paper, we propose a new statistical method, Model-based Association test Reflecting causal Status (MARS), that tries to find an association between variants in risk loci and a phenotype, considering the causal status of the variants. One of the main advantages of MARS is that it only requires the existing summary statistics to detect associated risk loci. Thus, MARS is applicable to any association study with summary statistics, even though individual level data is not available for the study. Utilizing extensive simulated data sets, we show that MARS increases the power of detecting true associated risk loci compared to previous approaches that consider multiple variants, while robustly controls the type I error. Applied to data of 44 tissues provided by the Genotype-Tissue Expression (GTEx) consortium, we show that MARS identifies more eGenes compared to previous approaches in most of the tissues; e.g. MARS identified 16% more eGenes than the ones reported by the GTEx consortium. Moreover, applied to Northern Finland Birth Cohort (NFBC) data, we demonstrate that MARS effectively identifies association loci with improved power (56% of more loci found by MARS) in GWAS studies compared to the standard association test.

## Introduction

Over the past decade, genome-wide association studies (GWAS) have successfully identified many variants significantly associated with diseases and complex traits. Unfortunately, those variants explain an extremely small proportion of phenotypic variation [1, 2] and there are many more variants with even smaller effects that we are yet to identify [1, 3–5]. Detecting all loci that harbor associated risk loci can help elucidate the biological mechanisms of diseases and complex traits. All biological follow up studies are performed on loci that harbor at least one significant variant. The standard association test used in GWAS examines one variant at a time to identify associated variants, we refer to this method as the univariate test.

It has been shown in previous works that many loci in the genome harbor more than one causal variant for a given disease or a trait [6–15]. In addition, recent works have demonstrated widespread allelic heterogeneity in expression quantitative trait loci (eQTLs) and complex traits [16, 17]. For a locus containing multiple causal variants with small effect sizes, the univariate test may be underpowered. Alternatively, an approach considering effects of multiple causal variants simultaneously may increase statistical power to detect signals for the locus by aggregating the effects of causal variants.

In this paper, we propose a new model-based method for identifying an association between multiple variants in a locus and a trait that we call Model-based Association test Reflecting causal Status (MARS). Our approach builds upon recent progress in fine mapping approaches that try to identify causal variants in a locus. Causal variants are the variants that are responsible for the association signal at a locus. However, at each locus, there are often tens to hundreds of variants tightly linked (linkage disequilibrium, LD) to the reported associated single nucleotide polymorphism (SNP), therefore, the LD hinders the identification of causal variants at the risk locus. CAVIAR [13] is one of the recent fine mapping approaches that estimates the probability of each variant being causal, allowing an arbitrary number of causal variants by jointly modeling association statistics at all the variants. We extend the likelihood model of CAVIAR to explicitly incorporate LD structure of data utilizing MVN distribution conditional on causal status of the variants. We perform a Likelihood Ratio Test (LRT) that computes the likelihood ratio of a null model, where none of the variants are causal, and an alternative model, where at least one of the variants is causal. For the significance test, we apply an efficient re-sampling approach.

Our method does not require individual level data, which is often not provided in GWAS. MARS only requires summary statistics and LD of variants in a locus, which can be obtained from a reference dataset such as HapMap [18, 19] or 1000 Genome project [20], and reports a *p*-value that indicates the significance of the association between the locus and the corresponding trait. This approach is related to set-based association tests that examine an association between a set of variants and a trait [21–23]. MARS outperforms these previous methods because the underlying model, building upon the model of CAVIAR, explicitly models the joint distribution of the observed statistics given multiple signals of associations. Furthermore, MARS uses a significance level which corresponds to the standard GWAS significance level facilitating interpretation.

Applied to several simulated data sets, we show that MARS robustly controls type I error and improves statistical power compared to the univariate test as well as the widely utilized set-based association test, fastBAT (a fast and flexible set-Based Association Test using GWAS summary data) [24]. Applied to data of 44 tissues provided by the Genotype-Tissue Expression (GTEx) consortium [25, 26], MARS identifies more eGenes, which are genes with at least one variant in cis significantly associated, than the ones reported by the GTEx consortium in most of the tissues, e.g. MARS identified 29% more eGenes than the ones reported by the consortium for the Whole Blood data and 57% of the extra eGenes, which were identified by MARS but not detected by the consortium, were reported in studies elsewhere. In order to demonstrate the increased power of MARS on real data, we followed a strategy of applying MARS on an older data set and validated additionally discovered loci using current datasets that have higher statistical power because they are much larger. Applied to the 2009 Northern Finland Birth Cohort (NFBC) data, we show that MARS effectively identifies more association loci than the univariate test and show that many of the new loci have since been discovered in recent GWAS studies.

## Results

### Overview of MARS

Causal variants are variants that are responsible for the association signal at a locus. The ultimate goal of the standard association test, which examines an association between each variant and a trait, is to find causal variants. We refer this method as the univariate test. However, often there exist multiple causal variants with small effect sizes in a locus. For these cases, the univariate test may not detect those associations due to its low statistical power. Alternatively, we can examine an aggregated effect of multiple variants simultaneously on the trait to increase statistical power. We developed a novel statistical method referred to as Model-based Association test Reflecting causal Status (MARS). MARS examines an association between a set of variants and a trait. MARS requires summary statistics estimated for variants (e.g. z-score) for a locus of interest and correlation structure, LD, between the variants, which can be readily obtained from a reference dataset. To test the association between a set of variants of a locus and a trait, MARS performs the Likelihood Ratio Test (LRT) to compute a test statistic, referred to as *LRT_score_*. We consider likelihoods of two models; the likelihood of the null model (*L*_0_) and the likelihood of the alternative model (*L*_1_). The null model assumes that there is no causal variant to the trait and the alternative model assumes that there is at least one causal variant to the trait. Then we compute the *LRT_score_* as *L*_1_/*L*_0_. Let’s say we are testing the association between *m* number of variants and a trait. Given the observed summary statistics we can compute the *LRT_score_* as follows:

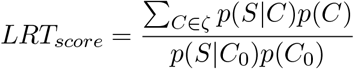

Here, *S* = [*s*_1_,⋯,*s_m_*]^**T**^ indicates summary statistics of *m* variants and *C* indicates the causal status of *m* variants. *C* is a binary vector of length *m*, where 0 indicates a variant is non-causal and 1 indicates a variant is causal. Specifically, *C*_0_ indicates the causal status where none of the variants are causal and *ζ* is a set that contains all the possible casual statuses except for the *C*_0_. Since there are *m* number of variants, there are 2^*m*^ possible causal statuses.

To assess an association significance for a locus, we utilize re-sampling approach, where we sample null statistics from a MVN distribution with corresponding LD and estimate *LRT_scores_* for the null statistics, to generate a null panel of 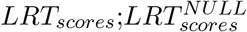. From the null panel, we estimate the siginificance of 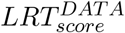, computed from the data. Figure 1 shows the basic overall process of MARS. The details and techniques to make this process computationally feasible for a big genomic data are described in the Materials and Methods.

**Figure 1:**
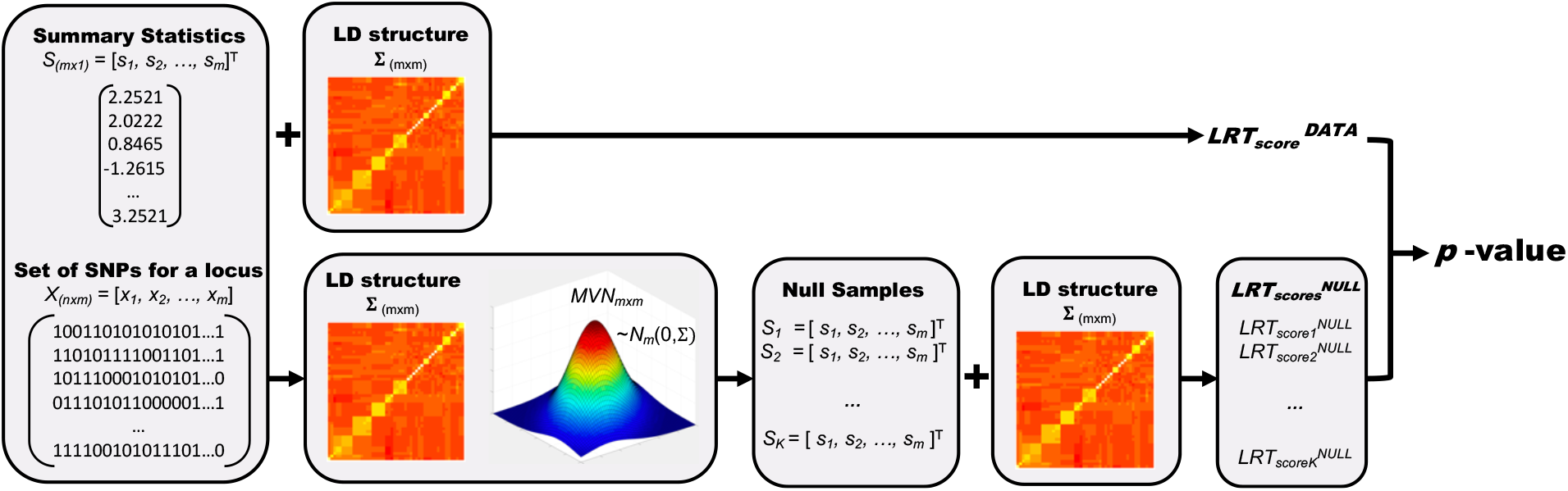
Overview of MARS. In this figure we assume that we are testing an association between a locus of m variants and a trait. The leftmost panel shows the input of MARS; *m* number of summary statistics for the locus and *n* × *m* matrix, containing genotypes of m SNPs for *n* samples. The next two panels in the bottom show the re-sampling process that we sample null statistics *K* times from a MVN distribution with a variance-covariance matrix of Σ, which contains LD of the genotypes *X*. The right most panel shows the process we estimate *LRT_scores_* for the null panel from which we compute a *p*-value of the data.

### MARS controls type I error while improves power in simulation studies

We demonstrate that MARS controls type I errors through simulations of null panels utilizing the GTEx data as a starting point and consider the SNPs ±lMb around transcription start, site (TSS) of 10 genes of Whole Blood data from the GTEx consortium [25, 26]. The half of the genes are randomly selected from genes reported as eGenes by the GTEx consortium and the other half of the genes are randomly selected from the rest of the genes, non-eGenes, of GTEx consortium [25, 26]. For each gene locus, we simulate 10^8^ number of null summary statistics according to the generative model described in the Materials and Methods, which uses the LD structure estimated from the genotypes of the SNPs in the locus and apply MARS to compute *LRT_scores_*.

To show that MARS controls type I error, the false positive rates are estimated for different thresholds of *α* = 0.01,0.05, and 0.1. The first half of the simulated data is used to compute a threshold of *LRT_scores_* for the corresponding *α; LRT_threshold_^α^* and the rest half of the simulated data is used to compute a quantile of *LRT_scores_* smaller than the *LRT_threshold_^α^*. Figure 2 (a) shows that MARS robustly controls type I error for all the examined gene loci as the false positive rates for different gene loci are very close to the corresponding *α* = 0.01, 0.05, and 0.1, accordingly.

To show that MARS increases statistical power, we perform extensive simulation studies for various scenarios and compare the power of MARS with those of the univariate test. Here, we define the univariate test as a set-based association test that uses the maximum summary statistic among the SNPs in a locus we are testing (for the details see Materials and Methods). The same gene loci from the previous section are used for the test. We estimate power of each gene locus for cases with 2 causal variants implanted with different effect sizes of λ = 4,4.5, 5, 5.5, and 6. For a fair comparison of the powers between the univariate test and MARS, we utilize the standard GWAS *p*-value threshold of 5 × 10^−8^. We simulate 10^8^ summary statistics under the null model of no effect to generate a null panel and 10^8^ summary statistics under the alternative model of effect size λ for 2 causal variants to examine the power. From the null panel, we find a LRT threshold that corresponds to the *p*-value threshold of 5 × 10^−8^. Then the power is estimated as a quantile of alternative cases that show *LRT_scores_* greater than the LRT threshold. For the detail, see Materials and Method. The percentage of power improvement is defined as (power of MARS - power of the univariate test)/(power of the univariate test) × 100. Figure 2 (b) shows that MARS increases statistical power compared to the univariate test. The extent of power improvements differs between the gene loci as the LD structures are different between the loci, but for all the cases, it is clear that the powers are improved over the univariate test. Depending on the effect size λ implanted in the simulated data, the power improves from 5.2% to 41.18% in our experiments and as expected, the smaller the effect size is, the better MARS performs over the univariate test. The results do not show noticeable differences between loci of eGenes and loci of non-eGenes used for the simulations.

**Figure 2:**
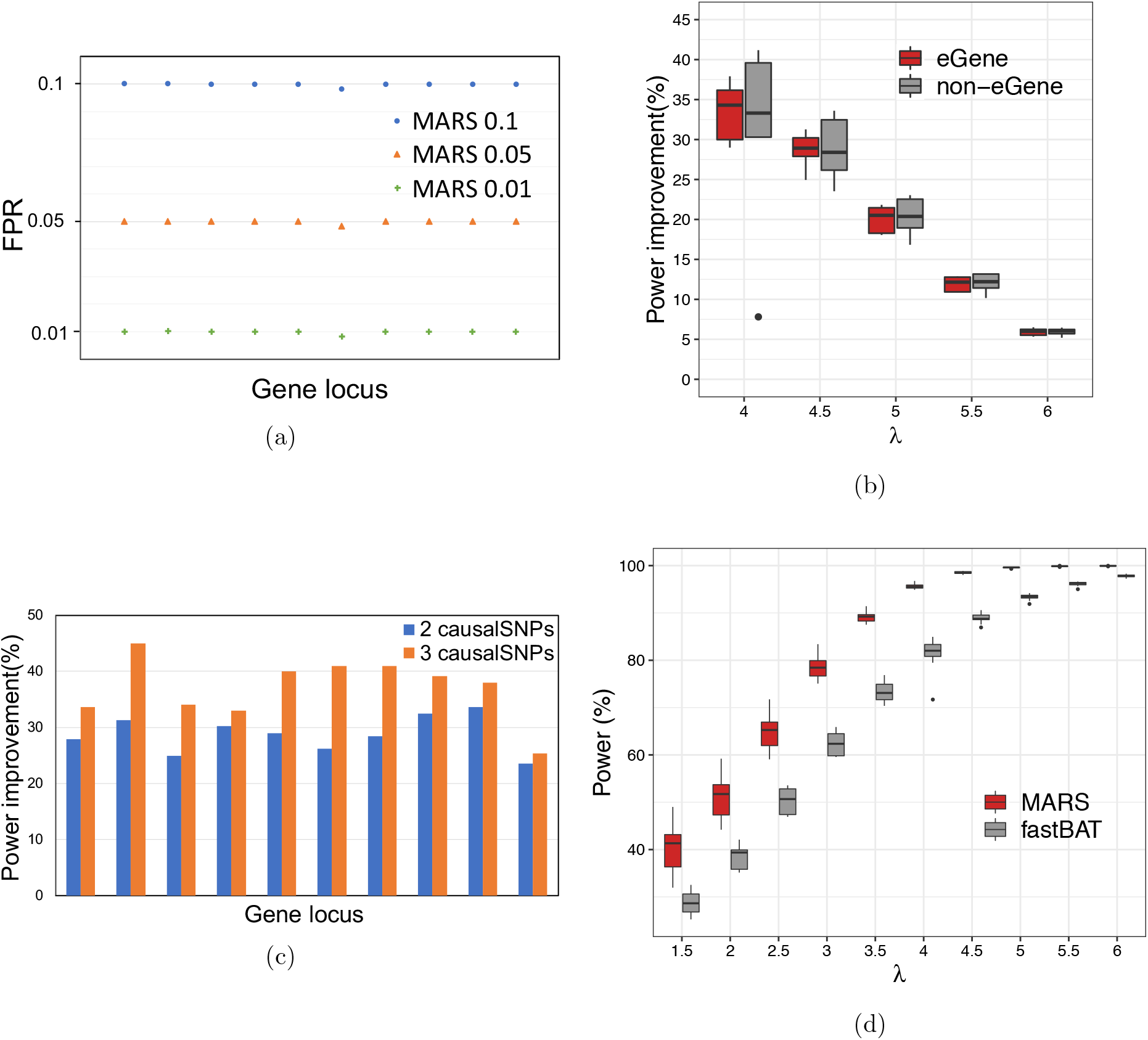
Comparison of eGenes identified by MARS and eGenes reported by GTEx consortium. (a) Plot, shows that MARS controls type I error. The x-axis shows 10 different gene loci used for the test, where the first 5 gene loci are from eGenes that are reported by the GTEx consortium. The y-axis shows the false positive rates. The green plus, orange triangle, and blue circle show the false positive rates at *α* = 0.01,0.05, and 0.1, accordingly, (b) A box plot, showing the percentage of power improvement, of MARS over the univariate test, for different, effect, sizes of 2 causal SNPs exist, in the data. The X-axis shows 5 different, effect, sizes of λ = 4, 4.5, 5, 5.5, and 6 used for the test. The Y-axis shows the percentage of power improvement. The red bars and the black bars show power improvement, when loci not. reported and reported as eGenes by the GTEx consortium are used for the test., accordingly, (c) A box plot, showing the power of MARS and fastBAT for different, effect, sizes. The X-axis shows effect, sizes of λ = 1.5,2, 2.5,3,3.5,4,4.5, 5, 5.5, and 6 used for the test. The Y-axis shows the power in percentage. The red bars and the black bars show power of MARS and fastBAT, accordingly.

In addition, we examine the cases, where 3 causal variants, each with an effect size of λ = 4.5, are implanted in the simulated data. As the number of causal variants increases from 2 to 3, MARS shows bigger power improvement over the univariate test Figure 2 (c). The result shows that the more the number of causal variants exist in a locus, the better MARS performs over the univariate test.

Besides the univariate test, we compared MARS with one of the widely used set-based association test methods, fastBAT [24]. Because of the heavy I/O of fastBAT, we used 10^5^ simulations and a threshold of 10^−5^, which is large enough to evaluate and compare the methods. We compute the power of MARS and fastBAT for different effect sizes of λ = 1.5,2,2.5, 3, 3.5,4,4.5, 5, 5.5,6 and show that MARS outperforms fastBAT for all the cases by improving power from 2% to 42% depending on the effect sizes in the experiments (Figure 2 (d)).

### MARS detects novel eGenes in GTEx data

Recently, a larger number of expression quantitative trait loci (eQTLs) studies have been reported. In particular, numerous *cis*-eQTLs, which are eQTLs that map to the approximate location of their gene-of-origin, have been identified. As part of this effort, GTEx consortium reported eGenes, which are genes with at least one *cis*-eQTL. We applied MARS to GTEx data to show that MARS can detect more eGenes than the ones reported by the GTEx consortium. Among the 44 tissues provided by the GTEx consortium, we first applied MARS to the Whole Blood data for the evaluation as the data contains the largest number of samples among all the tissues. To compare our results with GTEx’s results, we use 10000 number of simulations because it is the number of simulations used by GTEx consortium to compute ‘empirical *p*-values’ to select eGenes. To identify eGenes for MARS, we set the threshold as the border of empirical *p*-value between eGenes and genes other than those eGenes, referred to as non-eGenes, reported by the GTEx consortium. Figure 3 shows a Venn diagram comparing the identifications of eGenes by MARS and the ones reported by the GTEx consortium. MARS identifies 2043 extra eGenes that are not reported by the consortium, while MARS misses only 98 eGenes that are reported by the consortium [25, 26]. MARS and the GTEx consortium detects 6686 common eGenes.

**Figure 3:**
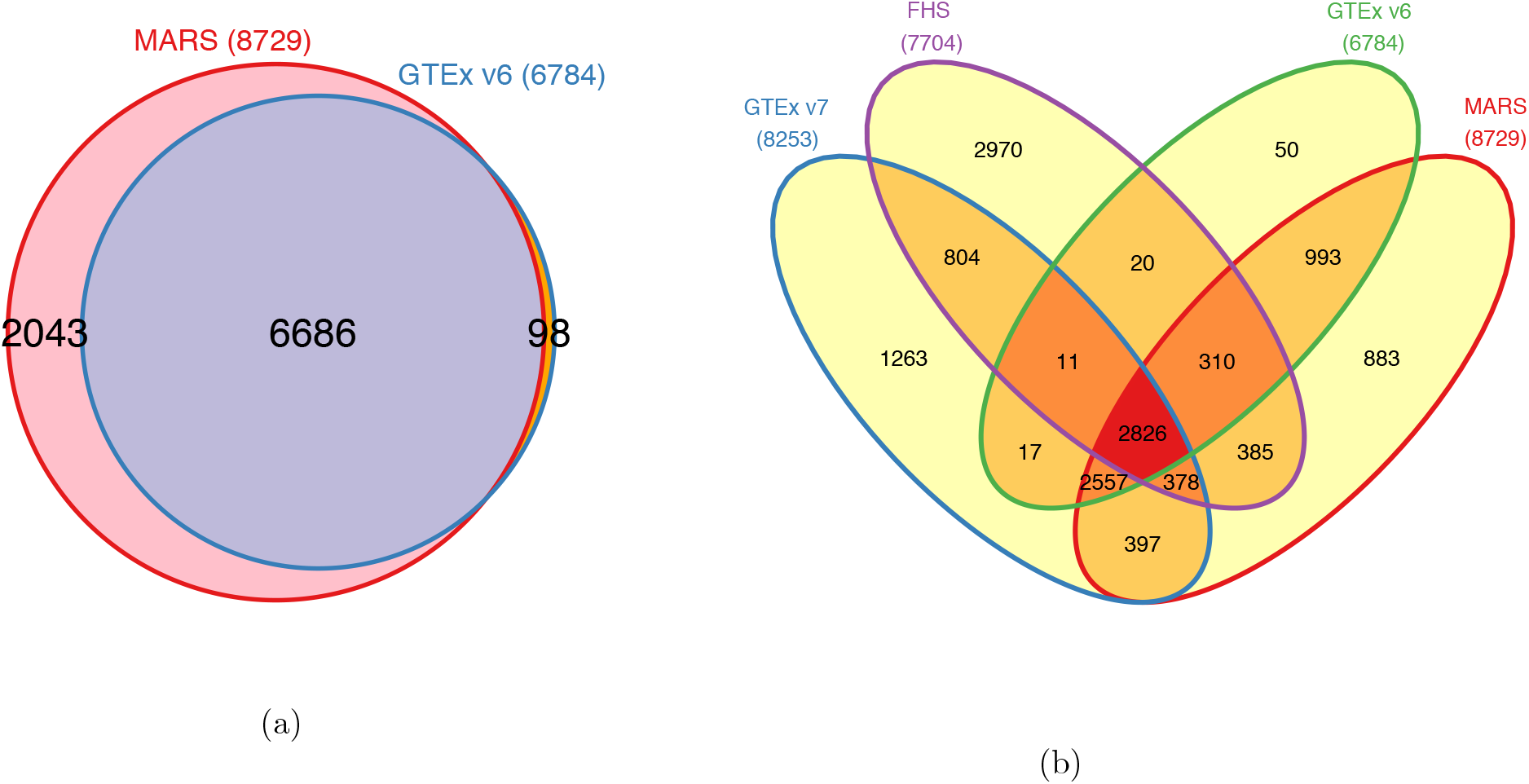
Comparison of eGenes identified by MARS and eGenes reported by GTEx consortium. (a) The red circle in the Venn diagram shows the eGenes identified by MARS and the blue circle in the Venn diagram shows the eGenes detected by GTEx version 6. Whole blood data is used for the analysis. (b) Venn diagram comparing eGenes identified by GTEx version 6, GTEx version 7, FHS, and MARS. Whole blood data is used for all three of the studies. The blue, purple, green, and red circle shows the eGenes identified by GTEx version 7, FHS (Framingham Heart Study), GTEx version 7, and MARS, accordingly. Note that MARS used data from GTEx version 6.

To verify that the eGenes identified by MARS are true associations, we compared those extra eGenes with those reported by other studies with larger sample sizes. Note that the results throughout the paper used data from GTEx version 6. Recently, GTEx version 7 has been published with more samples and improved technology in experiments. We expect more eGenes are detected in the newer version of the data as the power increases with the number of samples and etc. We compared the extra eGenes with the eGenes reported by GTEx version 7. In addition, we compared those extra eGenes with the eGenes reported by Framingham Heart Study (FHS) [27] that used the Whole Blood data on a large number of samples; 5257 samples. Figure 3b shows a Venn diagram comparing identifications of eGenes of four studies, GTEx version 6, GTEx version 7, FHS, and MARS. Among the 2043 extra eGenes, 57% of the genes (1160 genes) are reported in either GTEx version 7 or FHS; 763 genes are reported as eGenes in the GTEx version 7, 775 genes are reported as eGenes in FHS, and 378 genes are identified as eGenes by both of the studies, GTEx version 7 and FHS. Even with the older version of the data, MARS still finds more eGenes than GTEx version 7 and MARS is expected to identify even more eGenes if a data with larger number of samples is used in the further studies. Moreover, some of the genes among the 883 genes (Figure 3b) that are identified only by MARS but not by any other studies, GTEx version 6, GTEx version 7, nor FHS, have biological evidences of being eGenes based on many literatures. Variants of SP140 (ENSG00000079263) gene are known to be related to multiple sclerosis (MS) [28] and chronic lymphocytic leukemia [29]. Sille et al. have demonstrated that expression level of SP 140 is regulated by *cis*-eQTLs in lymphoblastoid cell lines [30]. Besides, the SP 140 protein levels are shown to be down-regulated by *cis*-acting mechanism in peripheral blood mononuclear cells (PBMCs) from MS patients [31]. HSPB8 (ENSG00000152137) has been recently identified as an eGene using PBMCs and the expression level of HSPB8 is known to be regulated by several SNPs [32]. Surfactant protein D encoded by SFTPD (ENSG00000133661) gene is known to be regulated by *cis*-acting manner in human blood [33], and CD83 (ENSG00000112149) has been recently identified as *cis*-eQTLs gene in CD19+ B lymphocyte [34]. Additionally, in Supplementary Information 2, we thoroughly analyzed randomly selected 100 genes and compared their *p*-values of MARS, the univariate test, the ones reported by the GTEx consoritium to show that MARS identifies more eGenes with better *p*-values. These results show that MARS is capable of identifying novel eGenes that cannot be detected with the standard association test approaches. The list of 2048 extra eGenes and their identifications in GTEx version 7 and FHS is provided in the Supplementary Information 2.

One of the advantages of MARS is that once the null panel of *LRT_scores_* for each gene has been established, it can be applied to the gene in any other tissues. Utilizing the null panel of *LRT_scores_* estimated from the Whole Blood data of the GTEx consortium, we computed *p*-values of the genes in all 44 tissues of GTEx using their summary statistics and LD structures. Figure 4 shows that MARS identifies comparable or more eGenes than the univariate test as well as the ones reported by the GTEx consortium in all the tissues. As expected, the number of eGenes identified by the univariate test and those reported by the GTEx consortium are very close to each other in all of the tissues. The numbers of genes are different between tissues due to the sample size differences and etc., and we used only the genes common in each tissue and the Whole Blood data of the GTEx consortium because the null panel of *LRT_scores_* was estimated for genes in the Whole Blood data. Comparison of eGenes identified by MARS, the univariate test, and the GTEx consortium for each tissue is provided in the Supplementary Information 3. In addition, we compared eGenes identified by MARS, GTEx version 6, and GTEx version 7 and provide Venn diagrams as the one in the Figure 3 for all of the tissues (Supplementary Information 4).

**Figure 4:**
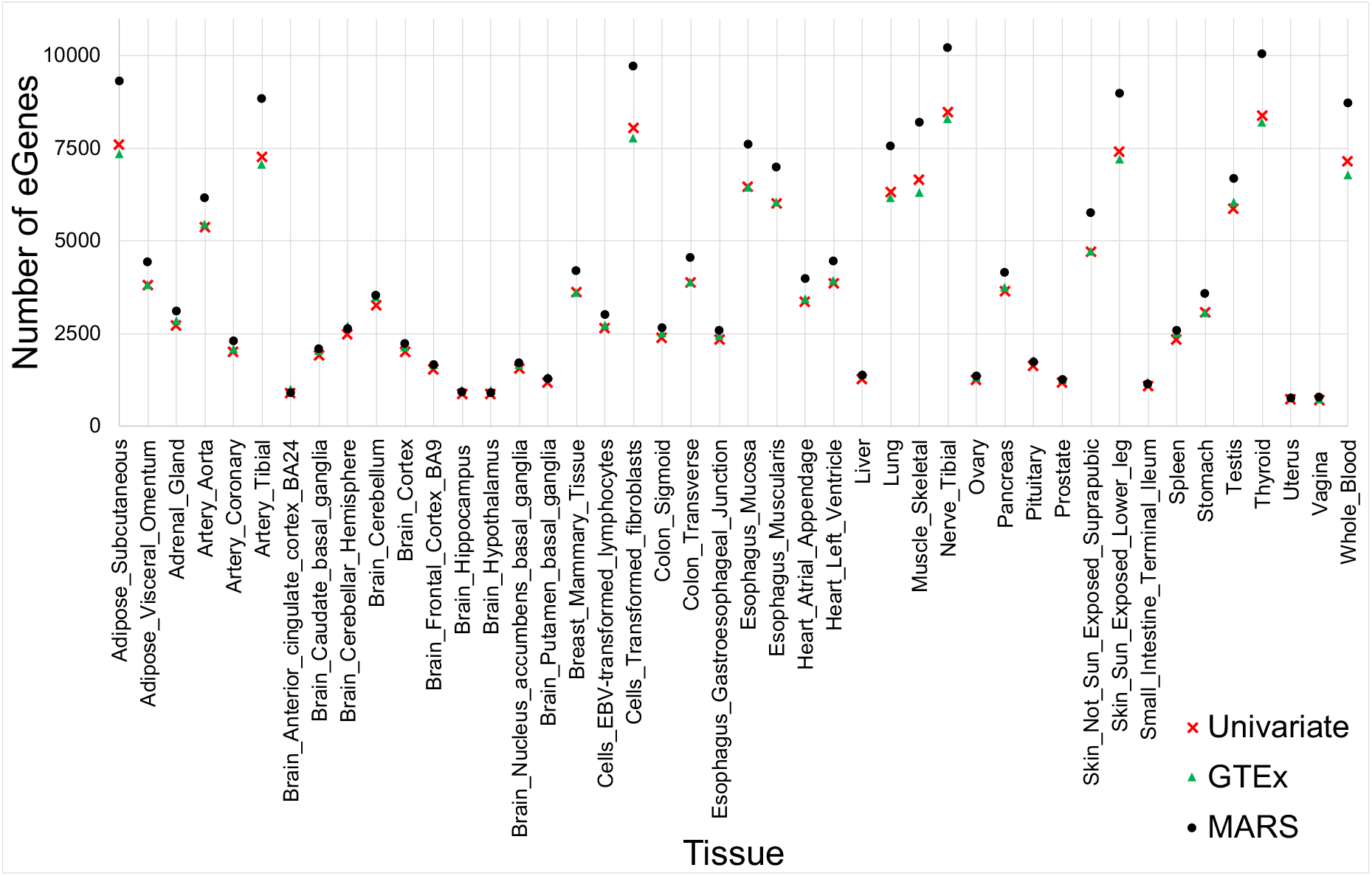
Number of eGenes identified by MARS, the univariate test, and those reported by the GTEx consortium. The x-axis shows the 44 tissues provided by the GTEx consortium and the y-axis shows the number of eGenes identified by each method. The black circle shows the number of eGenes identified by MARS, the red cross shows the number of eGenes identified by the univariate test, and the green triangle shows the number of eGenes reported by the GTEx consortium.

### MARS detects more set-based associations in GWAS

We show the effectiveness of our method on GWAS by applying MARS to the Northern Finland Birth Cohort (NFBC) data [35]. NFBC data consist of 10 traits collected from 5,327 individuals. The 10 traits are triglycerides (TG), high-density lipoproteins (HDL), low-density lipoproteins (LDL), glucose (GLU), insulin (INS), body mass index (BMI), C-reactive protein (CRP) as a measure of inflammation, systolic blood pressure (SBP), diastolic blood pressure (DBP), and height. For NFBC data we examined 51762 loci, where each locus is defined as ±1Mb of TSS of genes provided by the GTEx consortium.

To apply the standard *p*-value threshold of 5 × 10^−8^, MARS requires a lot, of sampling. To reduce the running time, we apply the idea, of importance sampling on MARS, which well approximates the *p*-value estimated from the original sampling approach, while reduces the number of sampling dramatically; from 10^8^ to 10^4^ (Supplementary Information 5). We refer to this efficient, version of MARS as fast,MARS. For the details, see Materials and Methods, section Fast, and space efficient, sampling for MARS. Figure 5 (a) shows that, for all the traits, fast,MARS identifies more or comparable loci that are likely to be significantly associated with the traits. Significantly associated loci identified by fast,MARS and the ones identified by the univariate test, are listed in the Supplementary Information 6 and Venn diagrams comparing the identifications of fast,MARS and the univariate test, are provided in the Supplementary Information 7. 471 number of loci are identified only by fast,MARS but, not, by the univariate test,. To verify those extra, loci, we searched the loci from other GWAS utilizing GWAS catalog [36]. As a result, several variants associated with 311 loci among the 471 extra loci were previously reported elsewhere in different studies [7, 37–61]. For example, rs6060369 locus associated with height was reported by large GWAS [44–46]. rs1800961 locus related to HDL was previously reported by large GWAS as well as meta-analysis GWAS [54–56, 60, 61]. rs6511720 locus related to LDL was discovered by several previous studies [40, 50, 51, 55, 56, 60]. A Venn diagram in Figure 5 (b) compares the number of loci identified by fastMARS and the univariate test as well as it shows the number of loci, for which at least one associated variant has been reported by GWAS catalog. The list of SNPs, the corresponding loci found by previous studies, and their detailed information including PubMed id and SNP position is provided in the Supplementary Information 8. Note that loci are defined based on the gene map of GTEx (±1Mb of TSS), thus some loci may overlap. For the details, see the Supplementary Information 8. These results demonstrate that fastMARS can efficiently identify novel associations in GWAS.

**Figure 5:**
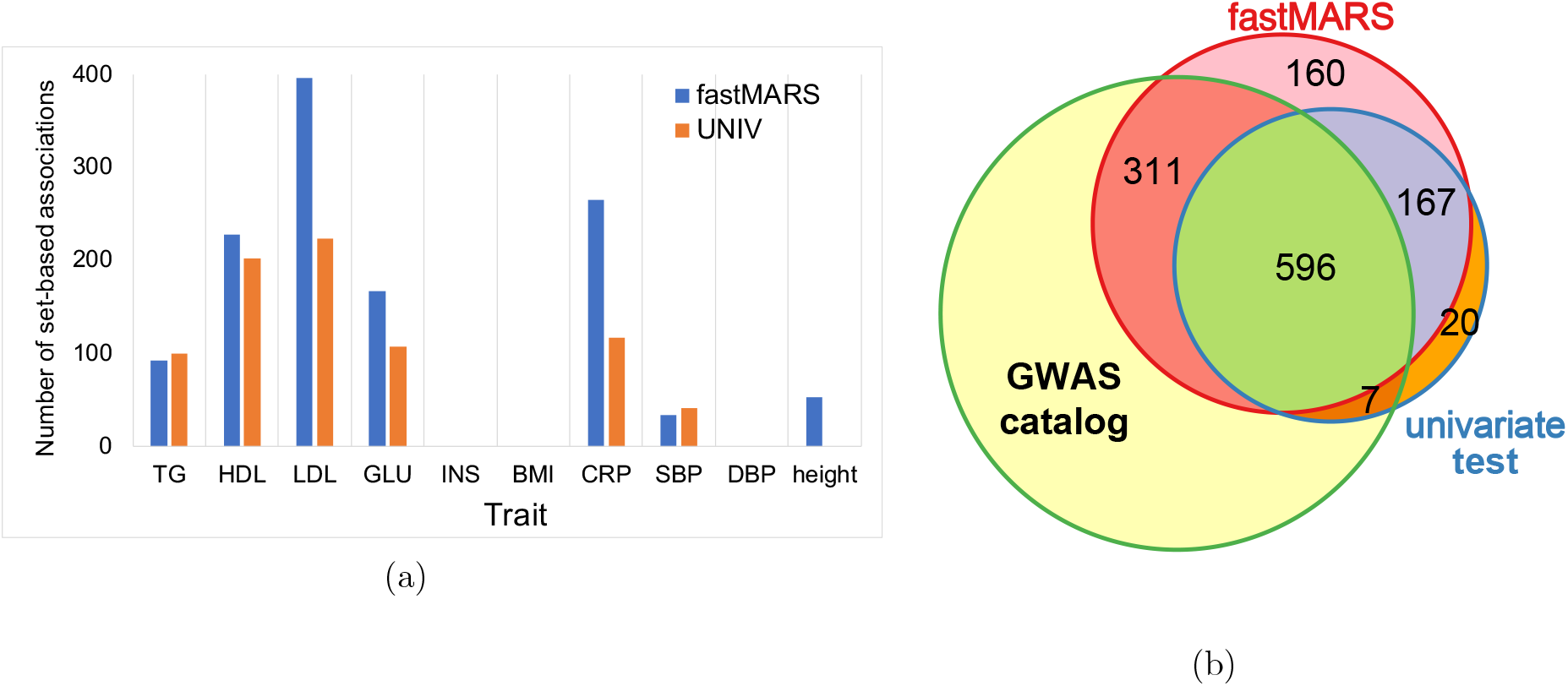
Significant associations identified by fastMARS and the univariate test in NFBC data. (a) Number of significant associations identified by fastMARS and the univariate test. The x-axis shows the 10 traits of NFBC data and the y-axis shows the number of set-based associations that is likely to be associated with the traits. Blue bars show associations identified by fastMARS and orange bars show associations identified the univariate test. The 10 phenotypes are triglycerides (TG), high-density lipoproteins (HDL), low-density lipoproteins (LDL), glucose (GLU), insulin (INS), body mass index (BMI), C-react.ive protein (CRP) as a measure of inflammation, systolic blood pressure (SBP), diastolic blood pressure (DBP), and height, (b) A Venn diagram showing the number of loci found by GWAS catalog for 10 traits. The red circle shows the number of loci identified by fastMARS, the blue circle shows the number of loci identified by the univariate test, and the green circle shows the number of identifications that were reported by GWAS catalog.

## Discussion

Great efforts have been spent on finding the hidden heritability and many studies suspect that single level variant test misses signals due to small effect sizes and power problems. An approach that examines multiple variants together may increase statistical power to detect risk loci with small effect sizes. Moreover, interpreting Genome-Wide Association Studies (GWAS) at a gene level is an important step towards understanding the molecular processes that lead to disease [62, 63]. Several statistical approaches have been proposed that test the association between a set of variants and a trait or disease status, however, they simply use naive statistics such as a mean or sum of *χ*^2^ of statistics in the risk loci [23, 24, 64, 65].

Our method examines an association between a set of variants and a trait considering causal status and LD between the variants, utilizing the model used in one of the recent fine mapping approaches [13, 66]. Our method uses summary statistics from each variant, therefore, does not require raw data which is often unavailable publicly. Another main advantage of our method is that, for a locus, once a null panel of test statistics has been established, it could be applied to the locus on other studies. For example, in eQTL studies, once null panels of genes have been established, we can readily access significances of the genes in other tissues, and the same strategy can be applied to different traits in GWAS.

We applied our method to extensive simulated data sets with different effect sizes and the number of causal variants to show our method improves power compared to previous approaches including a widely used set-based association test, fastBAT, while successfully controls the type I error. Especially, we show that when there are many causal variants with small effect sizes, our method performs superior over the standard univariate association test approach. Applied to Genotype-Tissue Expression (GTEx) data, our method identifies more or comparable eGenes than the standard univariate approach as well as the ones reported by the GTEx consortium in all of the tissues. In addition, using the Whole Blood data, we show that a large portion of the eGenes identified only by MARS was reported by other larger studies as well as some of them have biological evidence of being eGenes based on many literatures. Lastly, utilizing Northern Finland Birth Cohort (NFBC) data, we show the effectiveness of our method on the GWAS that our method effectively identifies more association loci in GWAS compare to the standard association test approach.

We note some limitations of our work. First, MARS is computationally costly compared to the standard GWAS method as MARS test significance of an association based on re-sampling approach. However, in practice, we introduce fast and space efficient sampling techniques including importance sampling to reduce the sampling time dramatically, which well approximates the original result, while we were able to successfully handle big eQTL data sets containing tens of thousands of genes or GWAS data sets with thousands of samples. Second, we limited the number of causal variants in a locus to reduce the running time in the experiments. It is a reasonable assumption as previous studies have reported that a relatively small number of causal variants exist in a region. There is a trade-off between the number of causal variants to be considered and the running time, thus, to detect loci with many causal variants and raise the accuracy, one can increase the number of allowed causal variants, which can be set as an option in the program. Third, MARS can only be applied to common variants but not to rare variants as the MVN assumption holds only for common variants. Lastly, MARS does not utilize existing functional data as some of current methods have utilized functional data to detect more eGenes [67–69]. We can extend the statistical framework of MARS to utilize functional data for the future work. Despite these limitations, MARS is a novel statistical method that can detect newly associated loci and increase the number of loci that we can perform follow up studies. Through which, MARS can increase our biological understanding of diseases and complex traits.

## Materials and Methods

### GWAS statistics

Consider GWAS on a quantitative trait where we genotype *n* individuals and collect a phenotype for the individuals. Let *X_i_* be a vector of length *n* with the standardized genotypic values (i.e., mean zero and variance one) of ith marker that we are testing and *Y* be a vector of length *n* with the phenotypic values. We assume that the data generating model follows the following linear additive model:

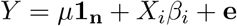

Here, *μ* is a mean of the phenotypic values, **1_n_** is a vector of *n* ones, *β_i_* is their coefficients, and **e** is a vector of length *n* sampled from 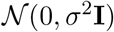 accounting for the residual errors, where **I** is an *n* × *n* identity matrix.

Under this model, the phenotype follows a MVN with a mean and variance as follows:

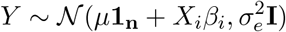

By maximizing likelihood of the model, we can estimate *β_i_* as follows:

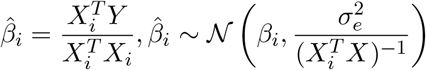

and the summary statistic is computed as follows:

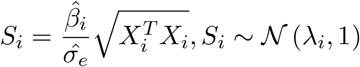

where λ_*i*_ is non-centrality parameter (NCP) and is equal to 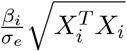. We obtain the estimated values for *μ*, **e**, and *σ_e_* as 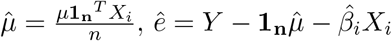, and 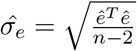.

### The effect of Linkage Disequilibrium on the Statistics

Consider the case that the *i*th SNP is causal to a phenotype and the *j*th SNP is non-causal but in LD with the *i*th SNP. The correlation between the two variants is *r*, which is approximated by 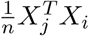. The effect size of the *j*th SNP is computed as follows:

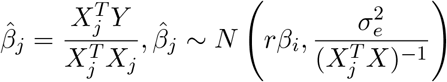

and the statistics for the *j*th SNP is computed as follows:

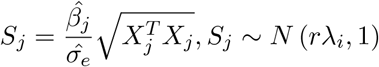

We can show that the covariance between the statistics is equal to the correlation of the genotypes as follow:

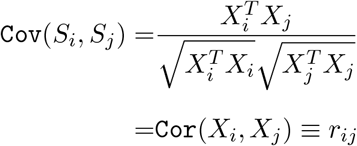

Then, the joint distribution of the summary statistics for the two variants given their NCPs, λ_*i*_ and λ_*j*_, follows a multivariate normal distribution as follow:

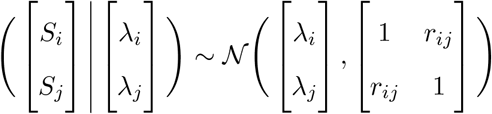

### CAVIAR Generative Model

Now we consider the case with *m* SNPs. Given the true effect sizes of *m* SNPs, Λ = [Λ_1_, Λ_2_, ⋯, Λ_*m*_], the summary statistics of *m* SNPs, *S* = [*S*_1_,⋯, *S_m_*]^*T*^, follows:

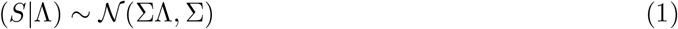

Here, Σ is a correlation matrix, where Σ{*i,j*} = *r_ij_*. We utilize Fisher’s polygenic model and assume that effect sizes follow a normal distribution. Let *C* be a binary vector of length *m* that indicates the causal status of *m* SNPs; 0 indicates a SNP is non-causal and 1 indicates a SNP is causal. Given a causal status *C*, we assume that the true effect size follows:

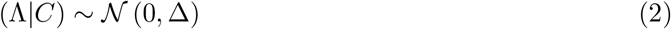

where Δ is a diagonal matrix with Δ{*i,i*} = *σ*^2^ if ith SNP is causal and Δ{*i, j*} = *ϵ*, otherwise. From equation (1) and equation (2), the likelihood of summary statistics follows a multivariate normal distribution as follows:

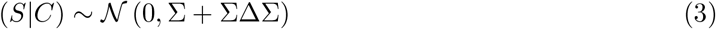

Then the likelihood function is given as follows:

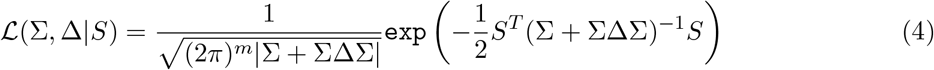

We use a simple model that a probability of a SNP begin causal is *γ*, which is independent from other SNPs. Therefore, we compute the prior probability as follows:

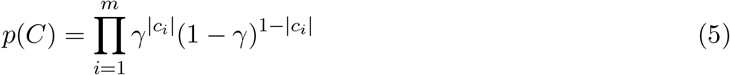

Here, |*c_i_*| = 1 if the *i*th SNP is causal and |*c_i_*| = 0, otherwise. Although we use a simple prior, we can incorporate external information by using SNP-specific prior *γ_i_*, which is the prior for the *i*th SNP, then the prior probability to a more general case is 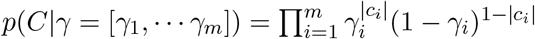.

### Model-based Association test Reflecting causal Status (MARS)

MARS examines an association between a set of SNPs and a phenotype of interest. For the test statistic, we utilize a likelihood ratio test (LRT). We consider likelihoods of two models; the likelihood of the null model (*L*_0_) and the likelihood of the alternative model (*L*_1_). The null model assumes that there is no causal SNP to the phenotype and the alternative model assumes that there is at least one causal SNP for the phenotype. Then we can compute the test statistic as *LRT_score_* = *L*_1_/*L*_0_. Given the observed marginal association statistics *S* and correlation matrix Σ, we can compute the *LRT_score_* as follows:

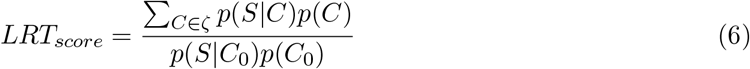

Here, we can compute the prior using equation (5) and the likelihood using equation (4). Since there are *m* SNPs, there are 2^*m*^ possible causal statuses. In practice, we limit the number of allowed causal SNPs to 2 or 3 as which is consistent with reports from previous studies that a relatively small number of causal SNPs exist in a region. In addition, as the size of genes are often very big, e.g. many genes contain more than 10000 SNPs within ±1Mb of TSS for the GTEx data, we order the SNPs by values of its summary statistics and used only top 50 SNPs for computing the *LRT_scores_* to reduce the running time and the space. Figure 6 (a) shows this practical implementation of MARS used for the experiments. This strategy reduces running time dramatically, while well approximates the results using all the SNPs in the loci (Supplementary Information 9) because the causal SNPs are expected to be included in the top 50 SNPs. In the case of limiting the number of causal SNPs up to 3 and using top 50 SNPs, there are 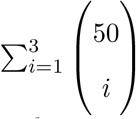 casual statuses to be considered and *ζ* becomes a set that contains all the possible casual statuses with 1, 2, or 3 causal SNPs.

**Figure 6:**
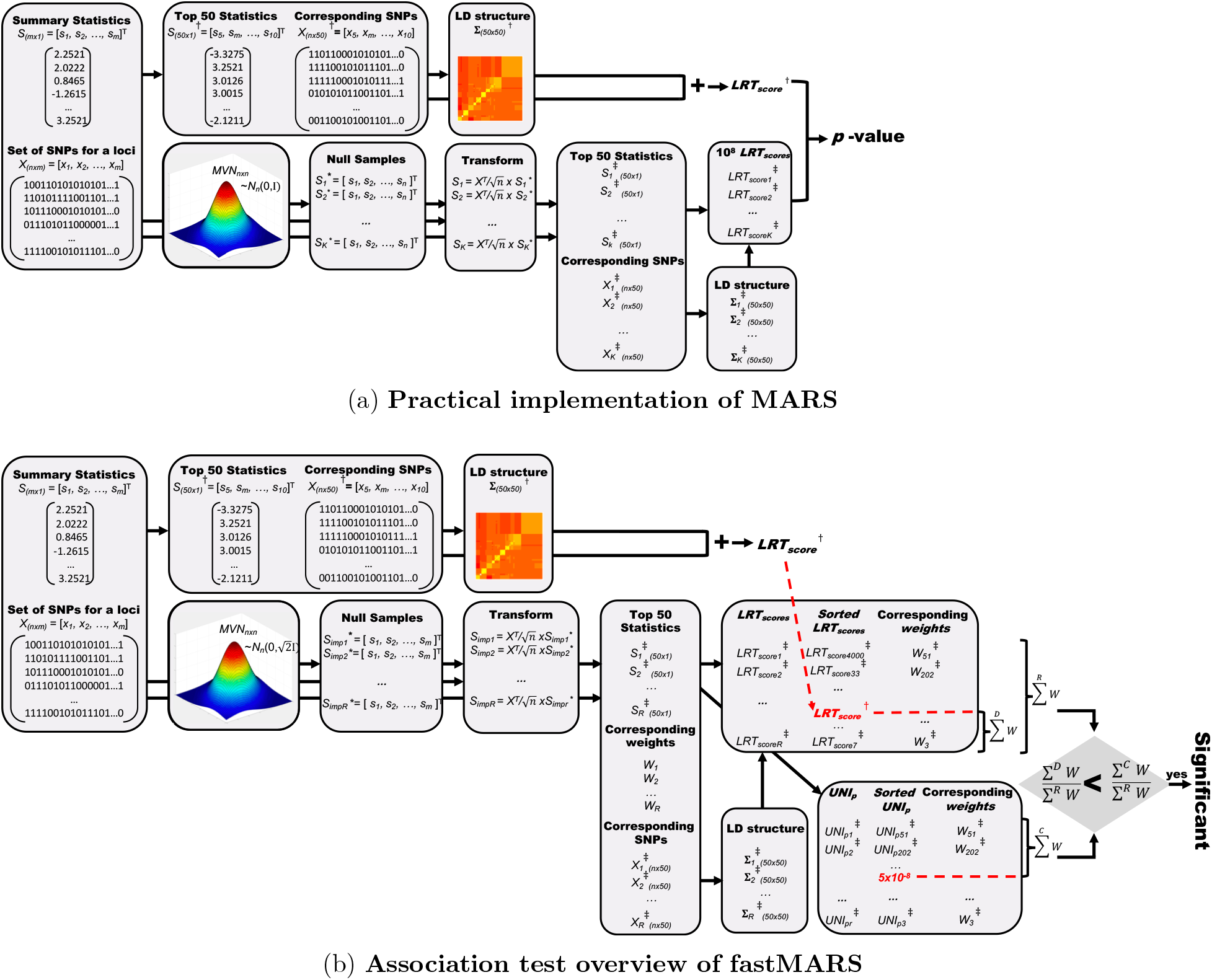
(a) To reduce the running time and the space, MARS uses top 50 statistics instead of using all the SNPs in the analysis in practice, (b) For GWAS, we introduce fast and efficient, sampling strategy.

### eGene detection in GTEx data

To identify an eGene, we examine the association between the gene expression levels and SNPs within ±1Mb of TSS of the gene, which can be the candidates of *cis*-eQTLs for the gene. To assess the significance for a gene, we sample summary statistics from a MVN distribution under the null hypothesis, *S* ~ *N*(0, Σ). Here, Σ is a variance-covariance matrix estimated from the SNPs within ±1Mb of TSS of the gene. We estimate the *LRT_scores_* for the null statistics to generate a null panel of 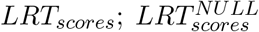, using the equation(6). Then, we order the SNPs by their values of summary statistics and select top 50 SNPs to compute the *LRT_score_* of the gene; 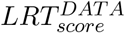, using the equation(6). The *p*-value of the gene is estimated as the quantile of 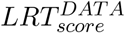 among 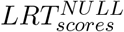 defined as follows:

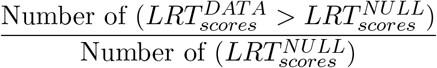

One of the advantages of MARS is that once the null panel, 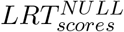, has been estimated for a locus, the panel can be applied to the locus in any other tissues or traits rapidly to compute a *p*-value. We use the Whole Blood data, which contains the most number of samples among the 44 tissues, to estimate the null panels of 23163 genes and applied the panels to all the other tissues. The *p*-value threshold to identify eGenes is defined as the border of empirical *p*-values of eGene and non-eGene reported by GTEx consortium, which is differ by tissues as GTEx used FDR approach to find their eGenes. The similar process has been applied for detecting eGenes in the univariate test except for using maximum summary statistic as its test statistic instead of *LRT_score_*.

### Power estimation

To show MARS increases statistical power over the univariate test, we compare the power between MARS and the univariate test. For a fair comparison, we utilized the standard GWAS *p*-value threshold of 5 × 10^−8^. We sample 10^8^ number of summary statistics under the null hypothesis, *S^NULL^* ~ *N*(0, Σ), as well as 10^8^ number of summary statistics under the alternative hypothesis, *S^ALT^* ~ *N*(ΣΛ, Σ). Here, Λ is a vector of length *m*, where *m* is the number of SNPs, containing zeros except for the causal SNPs. For example, for a simulation, in which two SNPs, let’s say SNP 1 and SNP 2, with effect size λ is implanted in the data, Λ is [λ, λ, 0,…, 0]. We examined the power for cases with 2 causal or 3 causal SNPs implanted in the simulated data, where the causal SNPs are randomly selected for each simulation. Then, we compute the *p*-value of *S^NULL^* using the univariate test; UNI*p^NULL^*, and find the quantile *q*, where the *p*-value equals to the standard GWAS *p*-value threshold of 5 × 10^−8^ as follows:

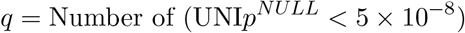

We compute the *LRT_scores_* of 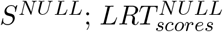, using MARS and set the *LRT_score_* at the quantile *q* as the threshold of 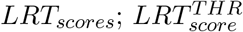, which satisfies the following equation:

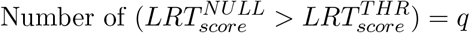

This 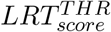 corresponds to the standard GWAS *p*-value threshold of 5 × 10^−8^. Now, we compute the *LRT_scores_* of 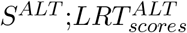, and the power of MARS is defined as the number of 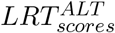 that is greater than the 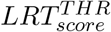 as follows:

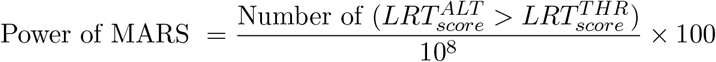

The power of univariate test is defined similarly by computing the *p*-value of *S^ALT^* using the univariate test; UNI*p^ALT^*, as follows:

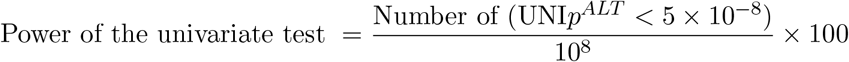

In the power comparison of MARS and fastBAT, the power estimation process is the same as the one described above except for that 10^5^ simulations and a threshold of 10^−5^ is used instead of 10^8^ simulations and the threshold of 5 × 10^−8^, accordingly. In the case, the univariate test is used to examine the quantile *q*, where the *p*-value of univariate test equals to 10^−5^. Then the quantile *q* is used to find a threshold of MARS and that of fastBAT to compute the power of each method.

### Fast and space efficient sampling for MARS

To access the significance of associations, MARS uses a re-sampling approach that requires a lot of sampling from MVN distribution. There are two main obstructions in this standard re-sampling approach. One is that a locus may contain many SNPs, for example, many genes in the GTEx data contain more than 10000 SNPs around ±1Mb of their TSS. When the number of SNPs *m* is very large, the standard re-sampling approach; *S* ~ *N*(0, Σ_*m×m*_), using the Cholesky decomposition [70] is impractical. This not only takes a lot of time but also a lot of space as Σ_*m×m*_ itself often takes a few gigabytes of space. We reduce the space and time complexity dramatically utilizing the fact that Σ_*m×m*_ is a covariance matrix of *X*; Σ_*m×m*_ = *X^T^X/n*, where *n* is the number of samples. Instead of sampling statistics from MVN with the variance-covariance matrix of Σ_*m×m*_; *S* ~ *N*(0, Σ_*m×m*_), we sample statistics from MVN with the variance-covariance matrix of *I_n×n_; S*\* ~ *N*(0, *I_n×n_*). This neither takes time nor space because in general *n* ≪ *m* and *n* is not big. Then we multiply *S*\* with 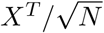 to compute the statistics 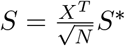.

The other main obstruction of the standard re-sampling approach is that the number of sampling required to find a proper threshold for MARS may be very big. For the GTEx data, we compared the eGenes with those reported by the GTEx consortium and 10000 sampling was performed as which is the number of samples used for computing their empirical *p*-values. However, for the GWAS analysis, MARS needs to perform a lot of sampling to find a LRT threshold that corresponds to the standard GWAS *p*-value threshold of 5 × 10^−8^. For the case, we apply importance sampling as follows. Instead of sampling from MVN with the variance-covariance matrix of *I*_*n×n*_; *S*\* ~ *N*(0, *I_n×n_*), we sample statistics from MVN with the variance-covariance matrix of 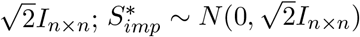). Then the new statistics from importance sampling becomes 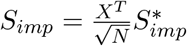. We record an additional information, referred to as importance weight, defined as follows:

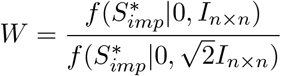

Here, *f* indicates the probability density function of MVN. We repeat the process of sampling statistics 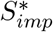 and computing *S_imp_* and *W, K* times. Let’s call each process as a run and after *K* runs we have a set of statistics {*S_imp1_, S_imp2_*,⋯, *S_impK_*} and a set of weights {*W*_1_, *W*_2_,⋯, *W_K_*}. Now we estimate an univariate *p*-value from each *S_imp_* and compute *p*-value threshold as the ratio of sum of weights that have the univariate *p*-value< 5 × 10^−8^ over the sum of all the weights as follows:

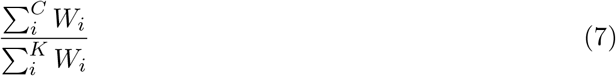

Here, *i* indicates the index of a run and *C* is a set containing indices of runs with the univariate *p*-value< 5 × 10^−8^. Given summary statistics of a locus, we access the significance of the locus by computing *LRT_score_* of the summary statistics; 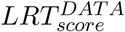. We compute the *K* number of *LRT_scores_* for the top 50 SNPs of the 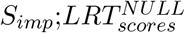, as well. Then we compute the *p*-value of the locus as the ratio of sum of weights that have 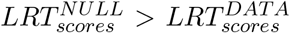 over the sum of all the weights as follows:

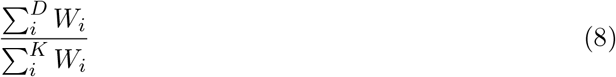

Here, *D* is a set containing indices of runs with 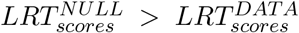. The association is significant if *p*-value estimated from the equation(8) is smaller than the *p*-value threshold estimated from the equation(7). Applied to randomly selected 10 genes, we find that the *p*-value estimated from 10^4^ number of importance sampling well approximates the *p*-values estimated from 10^8^ number of original re-sampling (Supplementary Information 5). Utilizing the importance sampling, we reduce the number of sampling dramatically from 10^8^ to 10^4^ on GWAS experiments. We refer to this fast and efficient version of MARS as fastMARS and Figure 6 shows the overview of association test for fastMARS (Figure 6 (b)).

For the GTEx data analysis, we used MARS as described in Figure 6 (a), where 10^4^ number of sampling performed and upto 2 causal variants considered. In the case, MARS took approximately 3.5 minutes to test a significance for an average sized gene with 7522 SNPs for 338 samples in our system. Using parallel processing, we were able to run the 23163 number of genes in several hours; approximately 3 hours for sampling and computing *LRT_scores_* and some extra times for pre-processing and post-processing the data. For the NFBC data analysis, which used 10^4^ original sampling, we used fastMARS as described in 6 (b), where 10^4^ number of importance sampling performed and upto 2 causal variants considered. In the case, fastMARS took approximately 50 minutes to test a significance for an average sized locus with 299 SNPs for 5326 samples in our system. Using parallel processing, we were able to run the 56319 number of genes in approximately 2 days.

### The standard univariate test and fastBAT

To compare MARS with a standard approach of set-based association test, we define an univariate test that uses a maximum summary statistic among the SNPs in the analysis locus. In addition, one of the widely used set-based association test, fastBAT (a fast and flexible set,-Based Association Test using GWAS summary data) [24] is used for the comparison. GCTA (genome-wide complex trait analysis) [71] program was downloaded from the GCTA website (http://gcta.freeforums.net/thread/309/gcta-fastbat-based-association-analysis) and ‘fastBAT’ option was used for running the GCTA-fastBAT.

### GTEx data

The summary statistics and genotypes of 44 tissues of GTEx data version 6 were downloaded from dbGap (https://www.ncbi.nlm.nih.gov/gap) and used to generate all the results throughout the paper. eGene list of GTEx data version 7 was download from dbGap as well and used only for the validation of eGenes that are identified by MARS applied on the GTEx data version 6. 23163 gene loci selected from the Whole Blood data was used for the analysis, which contain at least 50 SNPs in their ±1Mb of TSS. We generate null panel of *LRT_scores_* using the Whole Blood data as which contains the most number of samples, 338. The numbers of genes are different between tissues due to the sample size differences and etc., thus, for the eGene detection in 44 tissues, we used gene regions common in each tissue and the Whole Blood data.

### Northern Finland Birth Cohort dataset

The genotypes and 10 phenotype values of triglycerides (TG), high-density lipoproteins (HDL), low-density lipoproteins (LDL), glucose (GLU), insulin (INS), body mass index (BMI), C-reactive protein (CRP) as a measure of inflammation, systolic blood pressure (SBP), diastolic blood pressure (DBP), and height, over 5326 samples of the Northern Finland Birth Cohort (NFBC) dataset were downloaded from the dbGap. PLINK, a whole genome association analysis toolset (http://zzz.bwh.harvard.edu/pli is used to compute the statistics. For the set-based association test, gene map of the GTEx data, which contains 56319 gene positions, were used to define the loci to analyze. SNPs ±1Mb around transcription start site (TSS) of the genes were searched in the NFBC genotype data and 51762 regions with more than 50 SNPs were used for the analysis. 10^4^ number of importance sampling were performed to generate null panel to estimate *p*-values of MARS and the univariate test.

### Software availability and license

MARS software is implemented in C++ and the program, source code, installation and running instructions are available at http://genetics.cs.ucla.edu/MARS/. MARS is offered under the GNU Affero GPL, Version 3 (AGPL-3.0). For details of the license, see https://www.gnu.org/licenses/why-affero-gpl.html.

### Computing environment

For the experiments we used UCLA Shared Hoffman2 Cluster, which currently consists of 1,200+ 64-bit nodes and 13,340 cores, with an aggregate of over 50TB of memory. Note that the cluster is shared by many users and each user has a limitation of 100~500 running jobs in par-allel, depends on the memory and the time that each job uses. The details are provided at https://www.hoffman2.idre.ucla.edu.

### Ethics approval

No ethics approval was required for the study.

## Acknowlogement

This research was supported by Basic Science Research Program through the National Research Foundation of Korea(NRF) funded by the Ministry of Science, ICT & Future Planning, No. 2017R1C1B5017497 and No. 2018M3E3A1057752. E.E. is supported by National Science Foundation grants 0513612, 0731455, 0729049, 0916676, 1065276, 1302448, 1320589, 1331176, and 1815624, and National Institutes of Health grants K25-HL080079, U01-DA024417, P01-HL30568, P01-HL28481, R01-GM083198, R01-ES021801, R01-MH101782,NIH BD2K award, U54EB020403 and R01-ES022282. We acknowledge the support of the NINDS Informatics Center for Neurogenetics and Neurogenomics (P30 NS062691). F.H. is supported by NIH T32 DK110919 and F32HG009987.

